# Drug-loaded nanoparticles for cancer therapy: a high-throughput multicellular agent-based modeling study

**DOI:** 10.1101/2024.04.09.588498

**Authors:** Yafei Wang, Elmar Bucher, Heber Rocha, Vikram Jadhao, John Metzcar, Randy Heiland, Hermann B. Frieboes, Paul Macklin

**Affiliations:** Department of Intelligent Systems Engineering, Indiana University. Bloomington, Indiana 47408, USA; Department of Bioengineering, University of Louisville. Louisville, Kentucky 40292, USA

**Keywords:** agent-based modeling, anticancer drug-loaded nanoparticles, cancer nanotherapy, nanoparticle inheritance

## Abstract

Interactions between biological systems and engineered nanomaterials have become an important area of study due to the application of nanomaterials in medicine. In particular, the application of nanomaterials for cancer diagnosis or treatment presents a challenging opportunity due to the complex biology of this disease spanning multiple time and spatial scales. A system-level analysis would benefit from mathematical modeling and computational simulation to explore the interactions between anticancer drug-loaded nanoparticles (NPs), cells, and tissues, and the associated parameters driving this system and a patient’s overall response. Although a number of models have explored these interactions in the past, few have focused on simulating individual cell-NP interactions. This study develops a multicellular agent-based model of cancer nanotherapy that simulates NP internalization, drug release within the cell cytoplasm, “inheritance” of NPs by daughter cells at cell division, cell pharmacodynamic response to the intracellular drug, and overall drug effect on tumor dynamics. A large-scale parallel computational framework is used to investigate the impact of pharmacokinetic design parameters (NP internalization rate, NP decay rate, anticancer drug release rate) and therapeutic strategies (NP doses and injection frequency) on the tumor dynamics. In particular, through the exploration of NP “inheritance” at cell division, the results indicate that cancer treatment may be improved when NPs are inherited at cell division for *cytotoxic* chemotherapy. Moreover, smaller dosage of *cytostatic* chemotherapy may also improve inhibition of tumor growth when cell division is not completely inhibited.

This work suggests that slow delivery by “heritable” NPs can drive new dimensions of nanotherapy design for more sustained therapeutic response.

## 1. Introduction

There is an enormous interest in the application of nanomaterials in medicine for therapy and diagnosis, and understanding the associated nanomaterial-biological system interactions. Engineered nanomaterials have received particular attention due to the opportunities they offer with tailorable functionalities (e.g., desired shape, size, and surface compositions) to enable target-specific drug delivery with high efficiency and low side-effects [1, 2]. Sun et al. gave a detailed review of engineered nanoparticles for drug delivery in cancer therapy, covering different anticancer drugs, methods of controlled release, nanoparticles and drugs delivery, as well as study cases [3]. Shi et al. summarized recent work on cancer nanomedicine and novel engineering methods used to improve the understanding of tumor biology and nano-biological interactions to develop more effective nanotherapeutics for cancer treatment [4]. There are several advantages for nanoparticles working as therapeutic agents carriers in cancer therapy: (1) nanoparticles can increase therapeutic agents’ solubility and circulation half-life, and improve bio-distribution and permeability [3, 4]; (2) nanoparticles can increase targeting to tumor cells through selective binding to receptors overexpressed on the cells’ surface, by adding some targeting ligands (e.g., peptides, antibodies, and nucleic acids) on the NP surface, which would reduce collateral damage to healthy cells [5]; (3) multiple types of therapeutic agents can be delivered within the same NP, potentially reducing cancer drug-resistance by adjusting the ratio of different types of agents [6, 7]; (4) nanoparticles may improve cancer immunotherapy through development of synthetic vaccines by incorporating other molecules such as DNA, small interfering RNA (siRNA), messenger RNA (mRNA) and protein [4, 8]; (5) nanoparticles can be engineered to allow controlled release of encapsulated therapeutic agents based on particular physiological or external stimulus, e.g., pH, enzymes, temperature, or electromagnetic radiation [9].

Despite these advantages, NPs face a formidable journey from injection to effective intracellular action, including circulation in blood vasculature, uptake by immune cells, extravasation from capillaries, accumulation at the desired location, diffusion through extracellular space, endocytosis by cells, endosomal escape, intracellular localization and final action [10, 11], navigating abnormal physical and physiological properties that present delivery barriers [12]. Dogra et al. [9] and Stillman et al. [11] discussed three broad phases of NP transport from injection site to action site (tumor cells): vascular, transvascular, and interstitial, and NP-receptor binding, endocytosis as well as intracellular NP delivery. See recent reviews [13, 12, 14] for further background biology.

Mathematical modeling of cancer nanotherapy could be beneficial for exploring NP-loaded anti-cancer treatments affected by cell and tissue interactions. Simulations could help efficiently investigate nanomedicine design parameters to maximize cytotoxic effect while reducing systemic toxicity. Prior mathematical models of NPs traveling from injection site to action site have investigated NP transport through the vasculature [15], extravasation [16], tissue penetration [17, 18], endocytosis [19, 20, 21, 22] and intracellular trafficking. NPs may travel through different intracellular compartments, such as the cytoplasm, mitochondria, nucleus and lysosomes after internalization [14]. There is limited prior *in silico* modeling that addresses intracellular transport at this scale [11]. A detailed review of *in silico* modeling of cancer nanomedicine, across scales and transport barriers was presented in Stillman et al. [11].

Prior computational modeling of cancer nanotherapy typically used ordinary differential equations (ODEs), partial differential equations (PDEs) and/or coupled models of PDEs, and agent-based models (ABMs). For example, Dogra et al. used an ODE-based physiologically based pharmacokinetic (PBPK) model to investigate the whole-body NP pharmacokinetics and the impact of key parameters on delivery efficiency, such as NP degradation rate, NP size and tumor blood viscosity [23]. More recently, Dogra et al. developed an ODE-based pharmacokinetic/pharmacodynamic (PK/PD) model to perform translational modeling (from preclinical to clinical) for NP-mediated miRNA-22 therapy in triple negative breast cancer (TNBC) [24]. Such ODE-based PBPK, PK/PD and more complicated quantitative systems pharmacology (QSP) models have been widely used to quantitatively explore NP transport, delivery into tumor cells, and drug effects, allowing investigation of dose-response relation-ships and optimization of treatment frequency, important for evaluations of preclinical/clinical trails. These models generally cannot account for tumor cell heterogeneity and spatial interactions. Beyond ODE approaches, PDEs and/or hybrid PDE/ABM approaches have also been employed in cancer nanotherapy research. For example, Frieboes and co-workers evaluated cancer nanotherapy that included multiscale effects in a continuum tumor tissue representation, including NP transport, drug release and effects on vascularized tumor growth dynamics [25, 26, 27, 28, 29, 30]. These PDE and hybrid ABM/PDE models have advanced the modeling of tumor cell heterogeneity and spatial interactions (e.g., NPs and released-drug distribution within tumor tissue as well as tumor cell response to the microenvironments). However, since these prior models used PDEs to model the tumors, they could not track NP internalization and drug release for individual cells. In addition, because PDEs average space to smooth regions, they cannot consider the heterogeneity of cellular and subcellular/intracellular compartments, nor integrate cell-specific genetic or epigenetic effects [31]. PDE-based models generally neglect other key cell-cell interactions (e.g., physical pressure and biochemical signaling) and cell-extracellular matrix interactions, which are important determinants of cell phenotype that can affect therapeutic response.

In this work to model cancer nanotherapy, we develop an ABM framework and use it to investigate internalized NPs in individual cells, heterogeneity of tumor cells, and spatial interactions at the cell-NP, cell-cell, and cell-tissue scales. For the first time, we model in each individual cell a *population* of internalized NPs in a spectrum of drug release states. This allows us to more accurately track which NPs have been recently endocytosed by a cell (and thus have more bound drug to release), thereby giving a more nuanced view on drug release in the cytoplasm. In another aspect of this modeling, these NPs are “heritable”: when a cell divides, each daughter cell inherits half of the parent cell’s NP population. As we show in our model exploration below, heritable NPs admit new classes of therapeutic approaches, where drug-loaded NPs can persist through multiple cancer cell generations, potentially enabling long-term cancer control.

## 2. Methods

We use a hybrid discrete-continuum approach [32]: discrete cell agents model individual cancer cells, coupled with PDE representations of extracellular oxygen and NP concentration fields. In addition, each cell agent contains a system of ODEs to represent the population of internalized NPs across their drug release states, as well as the total amount of released drug.

### 2.1. Physicell: a multicellular simulation framework

In [33], Macklin and coworkers developed *PhysiCell* a framework for off-lattice agent-based simulations in multicellular systems biology, with a particular focus on cancer. In the framework, each cell agent has a phenotype with a hierarchically structured set of properties including state and rate parameters for cell cycling, cell death (apoptosis and necrosis), volume regulation (fluid and solid biomass, nuclear and cytoplasmic sub-volumes), motility, cell-cell mechanical interactions, secretion/uptake, and intracellular pathway reactions. More recent additions to PhysiCell’s built-in phenotype include cell transformations, phagocytosis, and effector cell attacks. Each cell agent can sample the microenvironment (through BioFVM [34], PhysiCell’s coupled diffusion solver), which is useful for modeling microenvironment-dependent triggers of standard cell processes. Each cell agent can have custom C++ rules assigned to model novel hypotheses (e.g., through rules of interpretable cell behavior [35]), which we use here to model NP internalization dynamics, NP drug release, anticancer drug effect on cancer cell growth dynamics, and distribution of NPs to daughter cells at cell division. See [33] for full algorithmic detail, numerical testing, and a variety of examples. *PhysiCell* has been applied to a broad variety of multicellular system problems, such as oncolytic virus therapy, cancer immunology, tissue mechanics, infection dynamics and tissue damage, cancer mRNA vaccine treatments, cancer cell migration, extracellular matrix remodeling, and cellular fusion, among others [36, 37, 38, 39, 40, 41, 42, 43, 44]. See [32] for detailed review of cell-based computational modeling in cancer biology.

We used PhysiCell [33] (version 1.7.1) to develop a multiscale ABM of cancer nanotherapy, and used it to investigate tumor cell growth and interactions with NPs encapsulating anti-cancer drugs. The model includes modules of NP internalization dynamics, drug release, NP “inheritance” and anti-cancer drugs effect on tumor cell phenotype. The overall approach is summarized in Fig. 1. The source code hosted on the GitHub repository is made public at: https://github.com/MathCancer/PhysiCell-nanobio.

**Figure 1:**
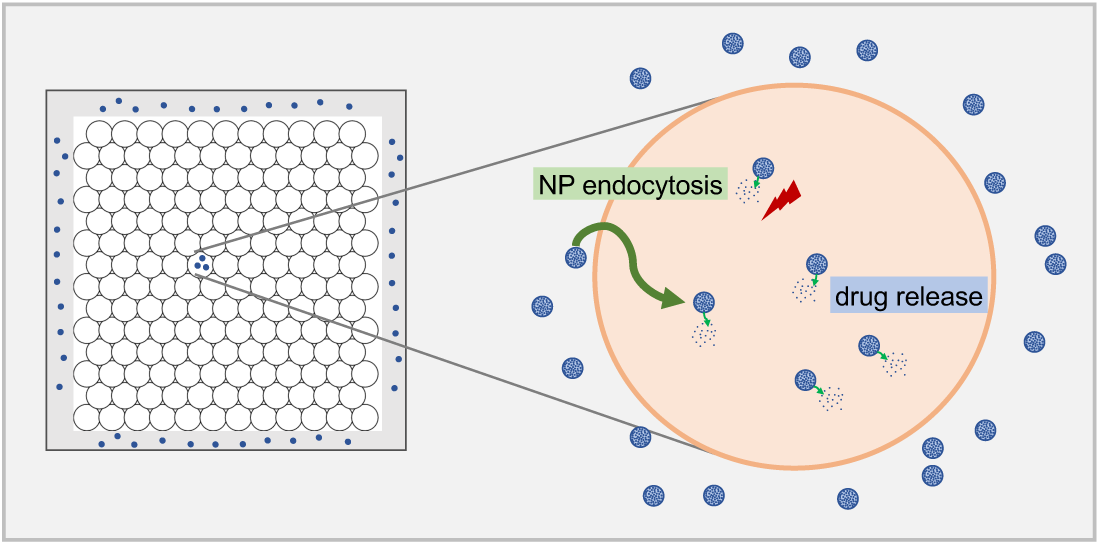
Schematic diagram of the overall mathematical modeling approach. A 2D tumor is numerically built, where NPs are released from domain boundary (“blood vessels”). Extracellular NPs are modeled as a continuum using PDEs to simulate diffusion through the microenvironment. After NPs are internalized inside of tumor cells via endocytosis, they start to release anticancer agents. Tumor cell phenotype (e.g., cycling, apoptosis, motility, mechanics and secretions) is impacted by the drug effects following rules defined in the model.

### 2.2. Cell-based model implementation details

We use the *Live* cell cycle model of *PhysiCell* [33, 45], where live cells can divide into two live cells with birth rate *b*. Each cell can divide, apoptose, or necrose based upon user-defined functions. We use the default built-in cell mechanics model (based on interaction potentials), and the built-in oxygen-dependent proliferation and necrosis: notably, the “Live” cell cycle transition rate (*b*) increases with oxygen availability above a minimal hypoxic threshold, and the necrotic death rate (*r*_nec_) increases below the threshold. See equations 1 and 2.

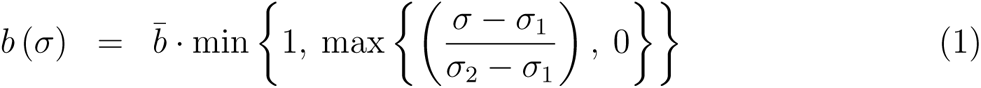

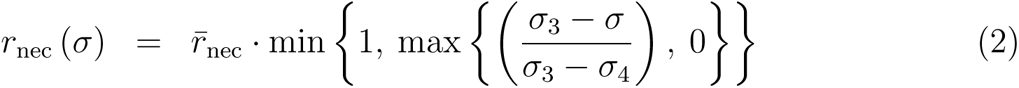

In equations 1 and 2:

- 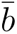: reference proliferation rate;
- *σ*_1_: oxygen proliferation threshold. Oxygen below which the proliferation ceases;
- *σ*_2_: oxygen proliferation saturation. Oxygen above which the proliferation rate is maximized;
- 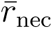: maximum necrosis rate;
- *σ*_3_: oxygen necrosis threshold. Oxygen value at which necrosis starts;
- *σ*_4_: oxygen necrosis maximum. Oxygen value at which necrosis rate reaches its maximum.

See *PhysiCell* [33] for further details and default parameter values, as well as recent experimentally-validated modeling [42] that used this functional form.

### 2.3. Oxygen and nanoparticle diffusion

We use BioFVM [34] to simulate diffusion of substrates in our model including oxygen, NPs, and cell-regulated signals. See equation 3. Specifically, if ***ρ*** is a vector of diffusible substrates with diffusion coefficients ***D***, decay rates ***λ***, bulk source rates ***S*** (which saturate at target densities ***ρ^∗^***), and bulk uptake rates ***U***, then problems of the following form are solved:

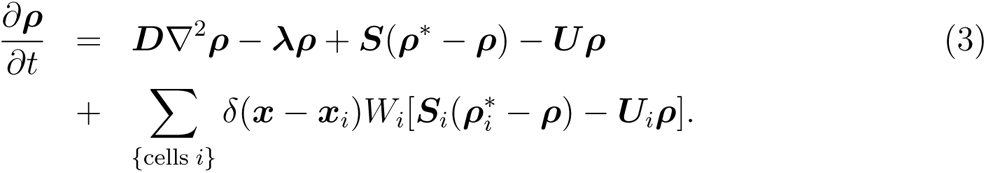

with Neumann (zero flux) boundary conditions. Here, for a collection of cells (indexed by *i*), with centers **x***_i_* and volumes *W_i_*, their secretion rates are **S***_i_* (with saturation densities 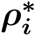) and their uptake rates are **U***_i_*.

See [34] for full details on the numerical method, implementation, and convergence testing. In [33], BioFVM was used as the default biotransport solver, which resulted in accurate and stable simulations, including for systems of moving cells.

In the model, oxygen and NPs are diffused from the boundary (to model a vascularized far-field condition) to the tumor center. Note that NP diffusion is slower than oxygen diffusion. Fig. 2 shows oxygen and NP diffusion contours.

**Figure 2:**
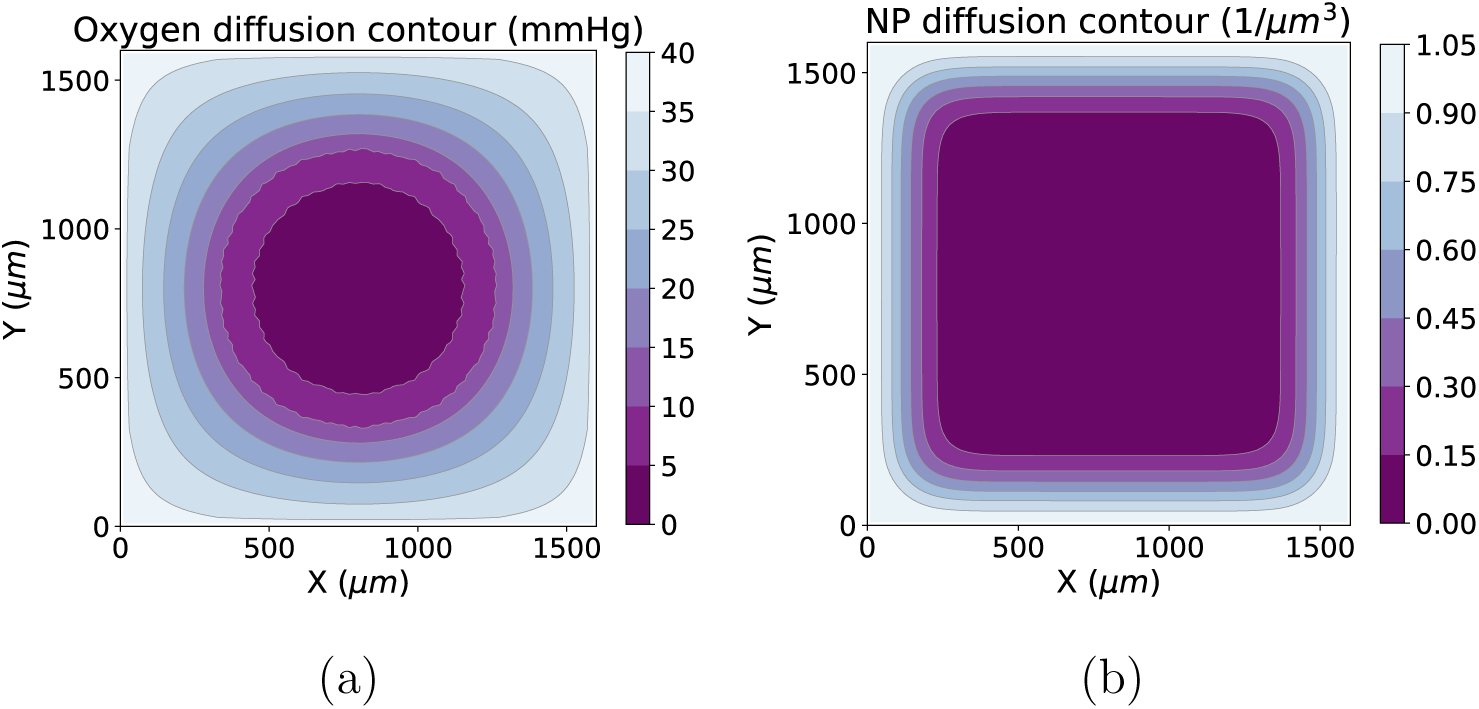
Oxygen (a) and NP diffusion (b) example results at time as of 24 hours (note that tumor cells have no cycling and death in the case)

### 2.4. Nanopaticle internalization dynamics

For the *cell-level* model, we propose a mathematical model of internalization dynamics based on experimental results reported in the literature. Based on observations in [46], the number of internalized NPs in a cell (*n_I_*) has a maximum, “saturated” value *n^∗^*, and the rate of internalization decreases as *n_I_ → n^∗^*. In addition, it was noted that the rate of internalization increases with the tissue concentration *ρ* of NPs (near and surrounding the cell). Accordingly:

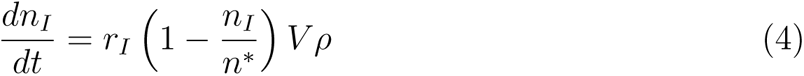

where *r_I_*is NP internalization rate, and *V* is cell volume.

For the *tissue-level* model, we adapt the conservation-based reaction-diffusion model from [34] the governing equation of NP transport:

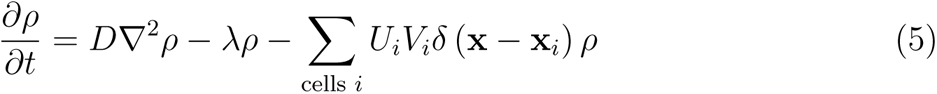

where *D* is NP diffusion coefficient, *λ* is NP decay rate, *U_i_*is the NP uptake rate for cell *i*, *V_i_*is cell *i* volume, *δ* (**x** *−* **x***_i_*) is Dirac delta function (equal to 1 inside cell *i* and 0 otherwise), and **x***_i_* is the cell *i* center.

The secretion of NPs is not simulated (or equivalently, only *net* uptake is modeled). We integrate over the domain Ω and simplify the result as follows (neglecting the effect of decay and diffusion by assuming short times):

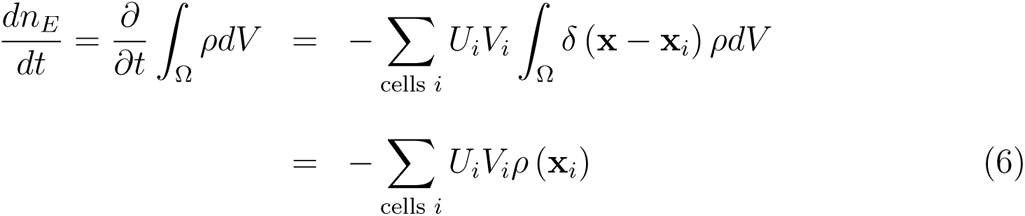

where *n_E_*is the total number of (non-internalized) nanoparticles within a tissue domain Ω.

*Matching the tissue-level and cell-level models*: Suppose we have a single cell, and match the tissue-level equation to the cell-level equation, neglecting the effect of decay by assuming short times. To maintain conservation of NPs, we require:

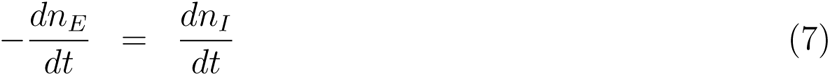

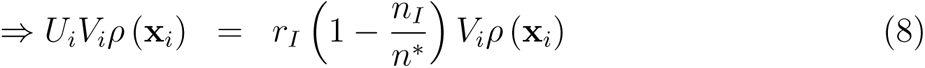

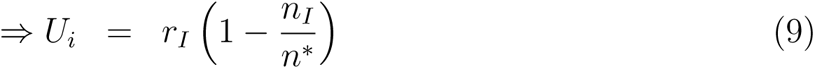

Because we can relate changes in the extracellular NP concentration (via uptake) to changes in intracellular number of NPs, we can track NP internalization for individual cells.

### 2.5. Nanopaticle drug release

Each NP arriving in a cell individually releases drug, leading to a *population* of internalized NPs with heterogeneous drug relase states. Therefore, we propose an “age”-structured model for drug release to model NP drug release states:

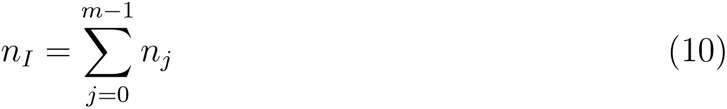

In Equation 10, *n_I_* is the total NPs (saturation) inside of each cell, *m* is the number of NP states, and *n_j_* is NP population in the *j_th_* state, with 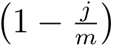 fraction of drug remaining in the NPs of in the *n_j_* compartment. We define the remaining drug in the NPs as follows:

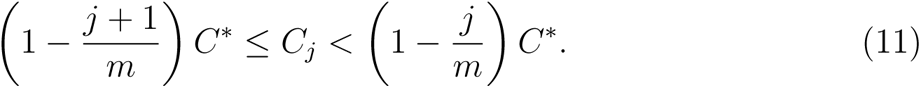

In Equation 11, *C_j_* is the remaining drug in the NPs within *j_th_* state, *C^∗^* is the initial drug in one NP. We can obtain the total released drug inside of individual cells:

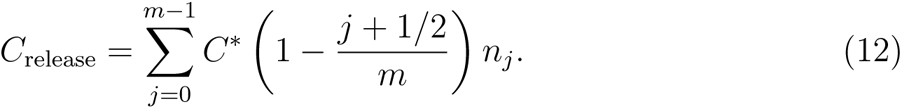

We assume the release rate in each state *r_j_* depends on the fraction of remaining drug in NPs (*C_r_*) and ratio of drug concentration over saturation inside of the cell:

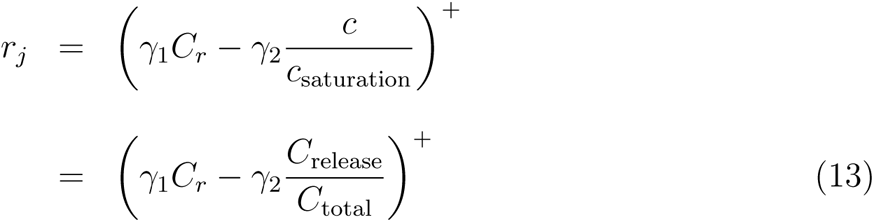

In Equation 13, *r_m−_*_1_ = 0 (in case of leak). 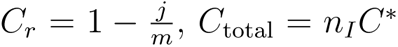. The function “(*x*)^+^” means non-negative part of *x*, i.e., (*x*)^+^ = max(*x,* 0). The ODE for drug release across the NP population is:

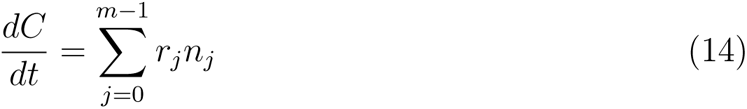

And the number of NPs in *j_th_* release state *C^∗^/m* is given by:

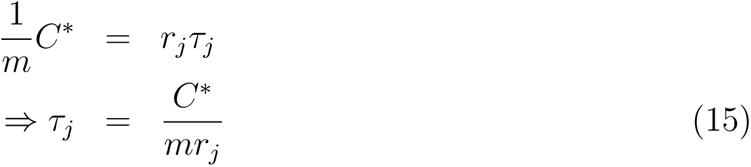

where *τ_j_*is the transit time of NPs in *j_th_*state. We summarize the population dynamics for NPs in *j_th_*state as following:

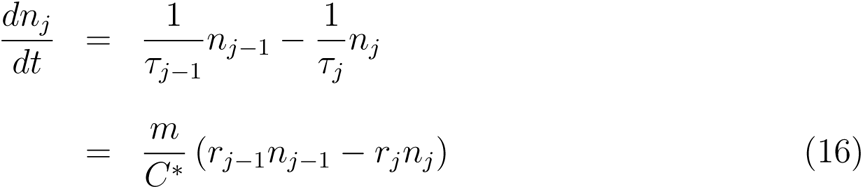

The boundary conditions for population dynamics in the first and last states (Equation 16) are:

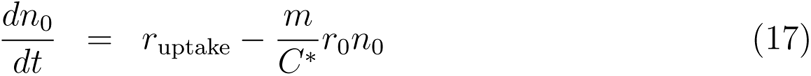

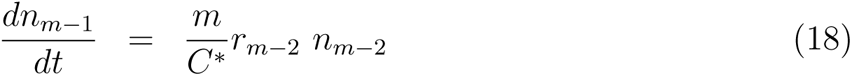

where *r*_uptake_ is NP endocytosed rate. We also assume all the drug would be released as long as NPs reached the last state.

See Fig. 3 for overall diagram of drug release. With the proposed model, we can track anti-cancer drug concentration and NP states inside individual cells. Fig. 4(b-d) present results of NP states in different time for the orange box area of Fig. 4(a). Tumor heterogeneity in NP internalization and drug release can be clearly observed.

**Figure 3:**
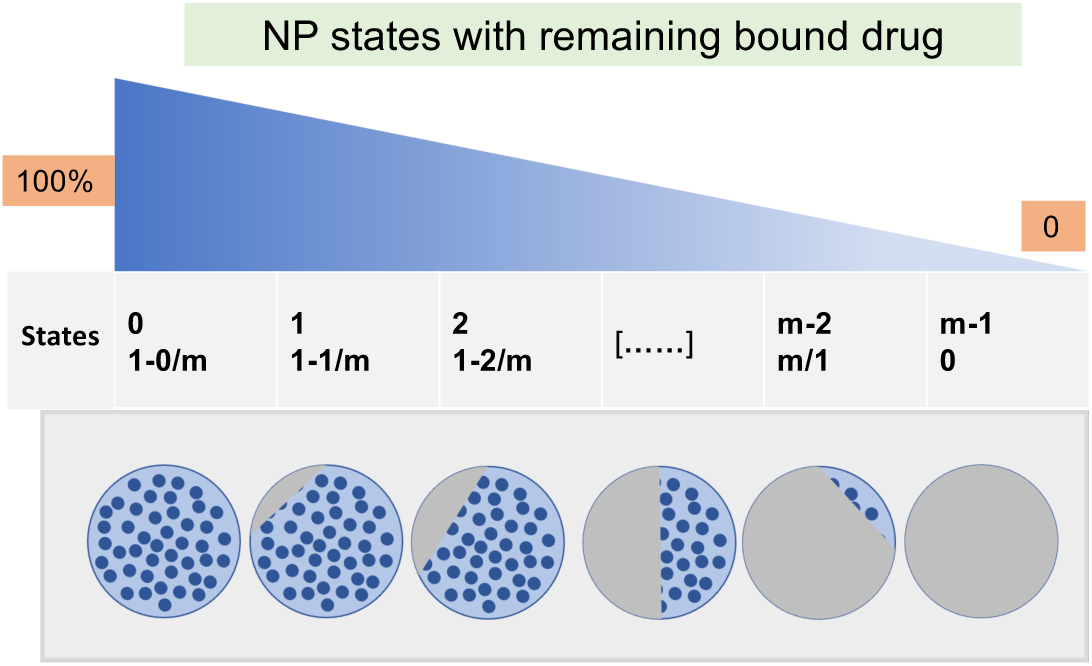
Schematic diagram of NP population dynamics in various states of drug release. In the model, each NP has *m* states (from 100% to 0), which means how much of drug remains in the NP.

**Figure 4:**
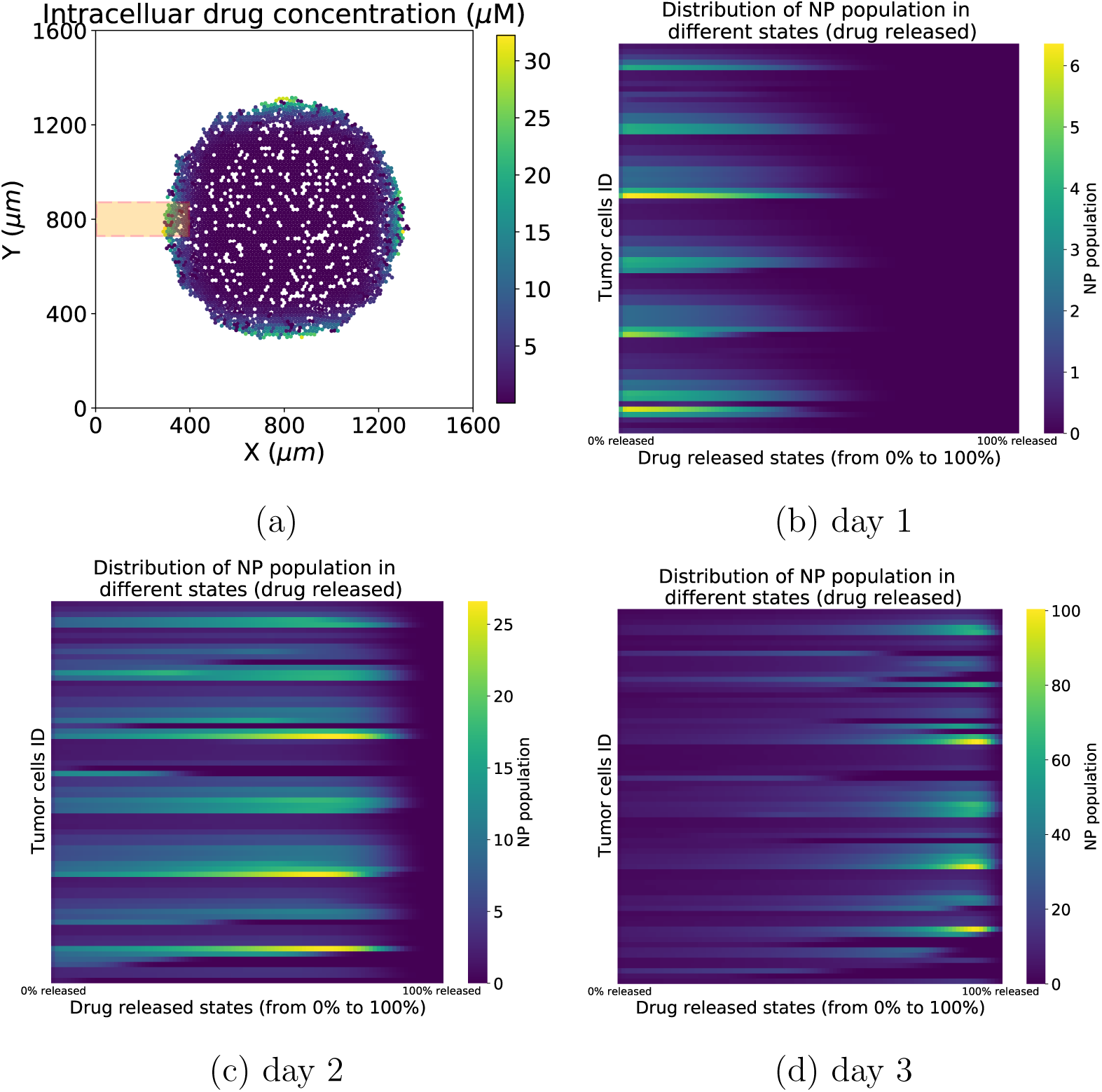
Distribution of NP drug release states, as tracked in individual cells. In (a), we plot the early intracellular drug concentration across the whole tumor, early in treatment. The orange box area of (a) (400 *µm ×* 150 *µm*) labels tumor cells analyzed in (b-d). In (b-d), we visualize the distribution of drug release states across the NNN labeled cells, (from 0 to 100%).

### 2.6. Nanopaticle “inheritance”

When a cell divides, conservation of mass dictates that the number of NPs in the daughter cells should equal those that were present in the parent cell; nonetheless, to the best knowledge of our knowledge, no other study to date has modeled and explored cancer nanotherapy where NPs are divided among the daughter cells at cell division, just as the cytoplasm containing the NPs. In order to simulate this effect, we aim to investigate how “inherited” NPs impact the treatment’s response. See Fig. 5 for model diagram. We will investigate in detail how NP “inheritance” impacts cancer treatments in Section 3.

**Figure 5:**
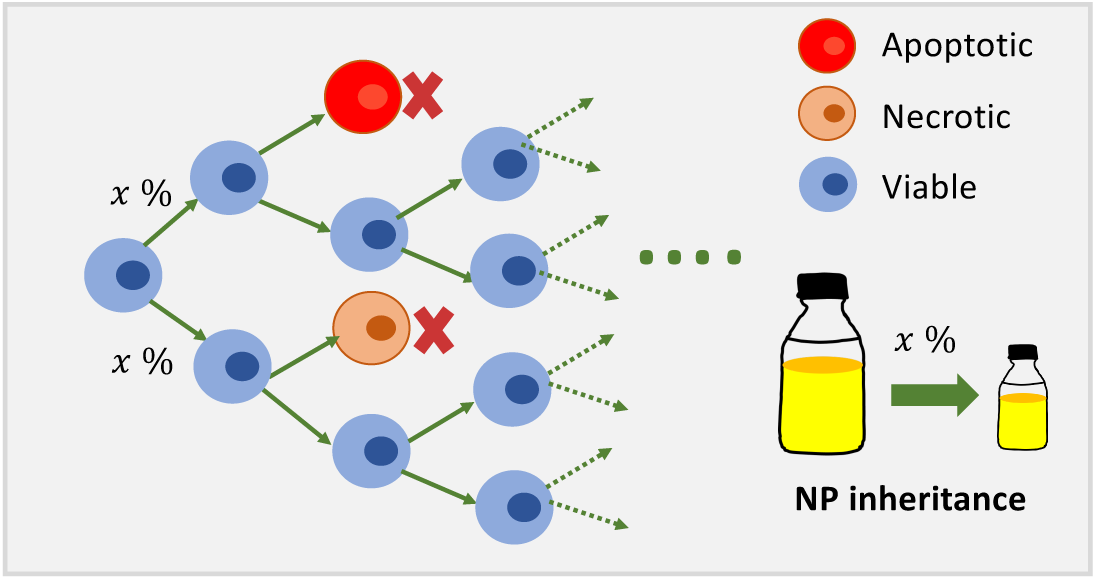
Schematic diagram of NP inheritance at cell division. In the model, tumor cells can divide, apoptose, or necrose. Two daughter cells can receive *x*% (0 *≤ x ≤* 50) of NPs from parent cell at cell division.

### 2.7. Anti-cancer drug effect on cell phenotype pharmacodynamics

We calculate drug effect based on intracellular drug concentration using sigmoidal (Hill) response functions. See applications of Hill functions in pharmacological modelling and multicellular system modeling in [47, 35]. Tumor cell phenotype (cycling or apoptosis rate, depending on the type of chemotherapeutic) is updated in the model by linearly interpolating the “base” phenotype (in the absence of drug) and maximal change in the phenotype under therapy, using the nonlinear drug effect *E* as the interpolating parameter. We use two common models for the drug effect *E*: Hill response functions, and area-under-the-curve (AUC) models.

*Hill functions model*: This “S-shaped” curve is given by

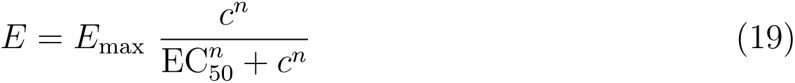

where EC_50_ represents that drug yields 50% of max effect; *E*_max_ means the maximum drug effect; *n* is Hill power.

*AUC model:* The drug concentration *c* may be replaced by an integration of time-*AUC* (Area Under the Curve) for some damage:

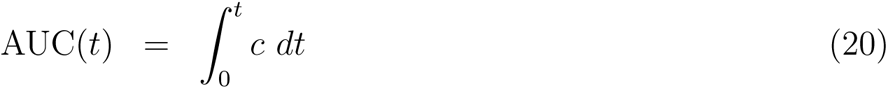

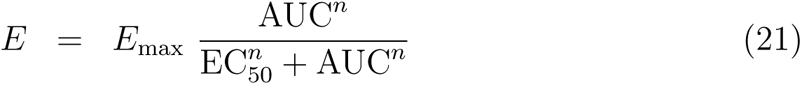

The drug may change cell phenotype (e.g., cycle, apoptosis, motility, mechanics, secretion) from a background rate (**r_0_**) to a max rate (**r**_max_):

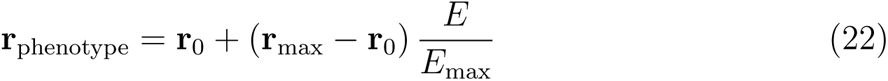

The model framework can also model other drug effects such as impact on cellular uptake rate, secretion rate, mechanics, or motility. Refer to our cloud-hosted app for full model demonstration.

### 2.8. Computing and cloud-hosted model app

Because the simulation model is stochastic, we ran 10 simulation replicates (each with a different random seed) for each parameter set. All simulation images are chosen from one representative replicate, and unless noted otherwise, all aggregate dynamical curves (e.g., viable cells) are reported as the mean and confidence interval (defined as *±* one standard deviation) over all replicates. Simulations were performed on the Big Red 3 supercomputer at Indiana University. See Fig. 6 for schematic diagram of large-scale parameters exploration for ABM framework). Each simulation was run on a single node using 24 threads. All jobs were submitted as a batch.

**Figure 6:**
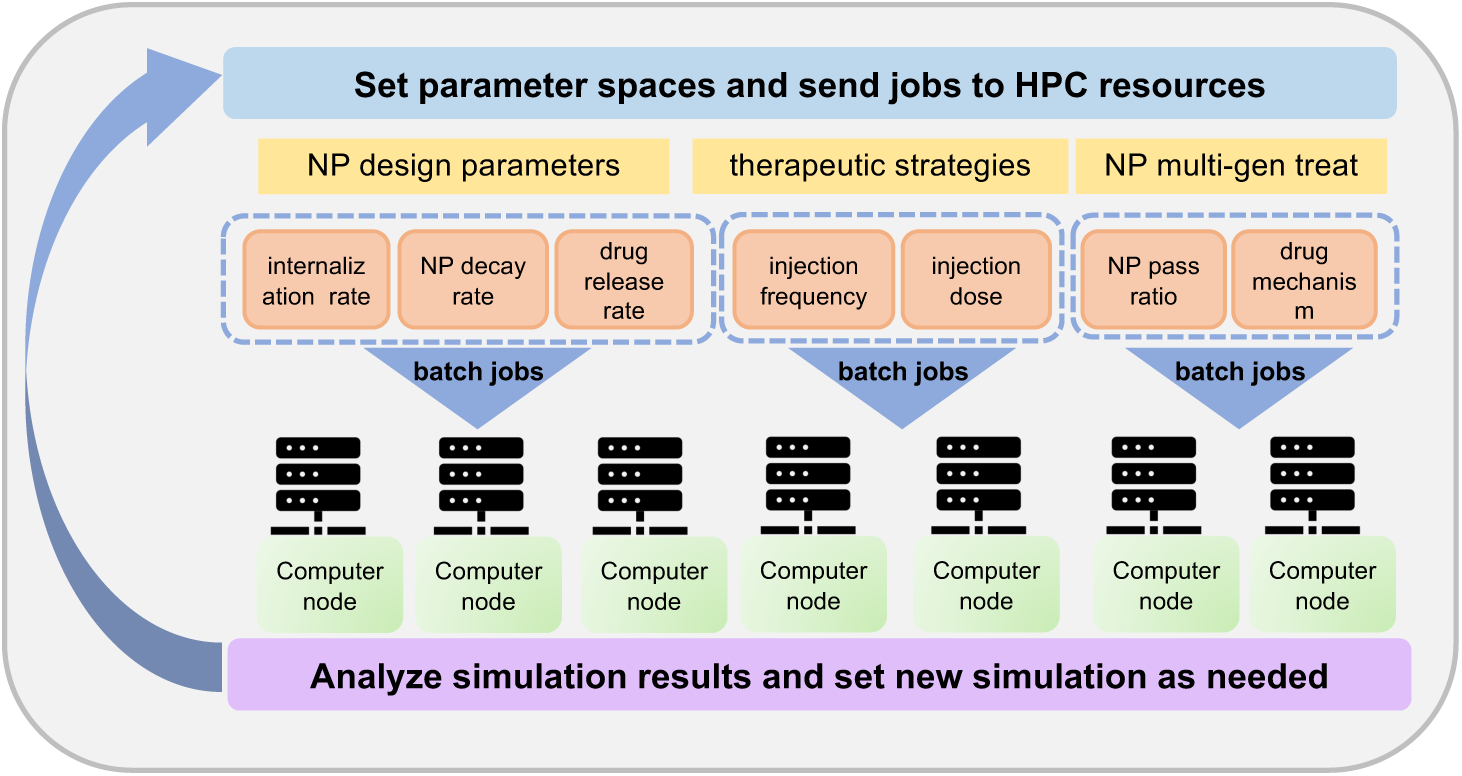
Schematic diagram of large-scale parameter exploration. In the investigation, parameter spaces are set and batch jobs are submitted to HPC for computing, collecting data and analyzing results.

We used *xml2jupyter* [48] to create a cloud-hosted version of this model: *pc4nanobio*; the model can be run interactively in a web browser at: https://nanohub.org/ resources/pc4nanobio [49]. Fig. 7 gives representative snapshots of the online model.

**Figure 7:**
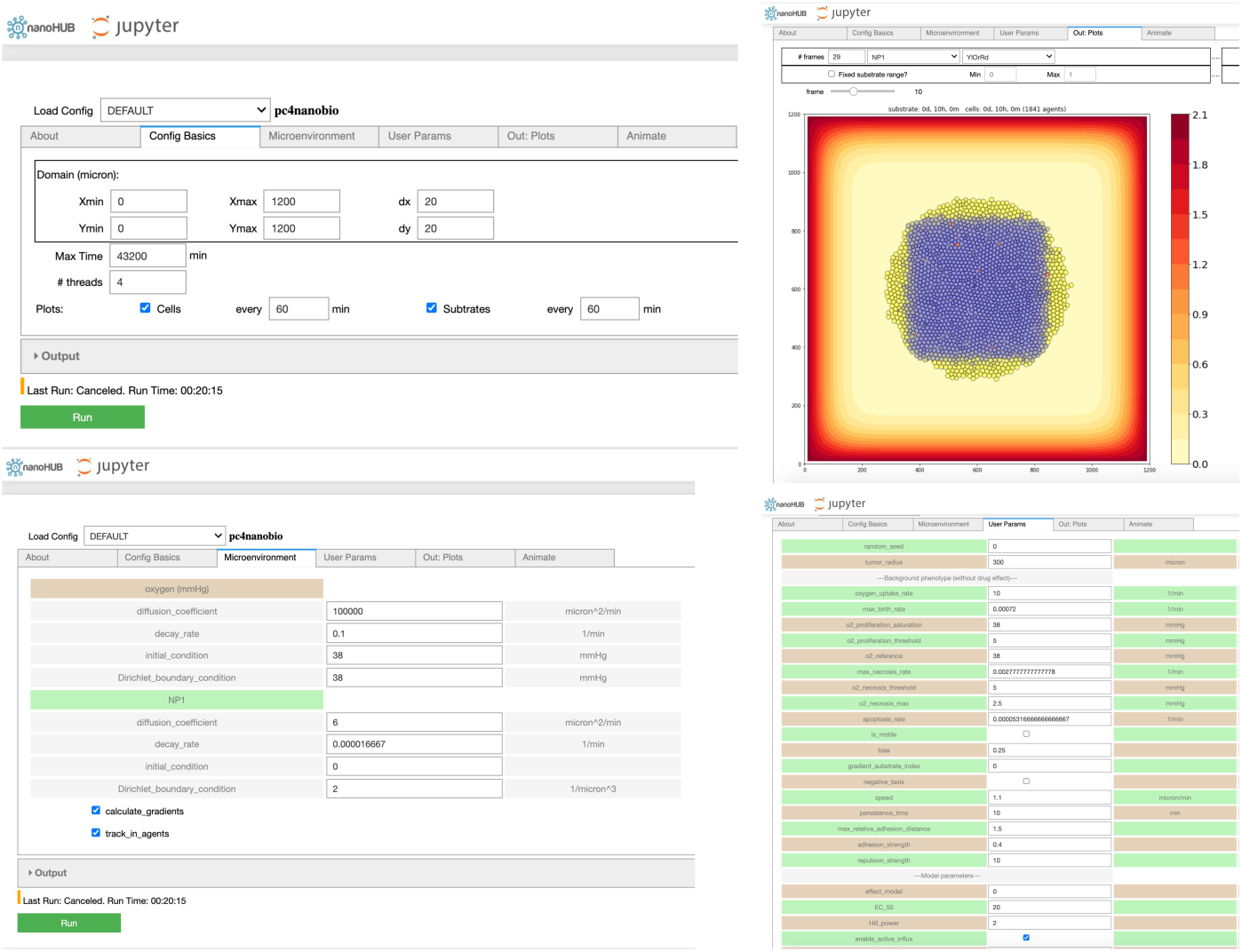
Cloud-hosted interactive model-*pc4nanobio* (version 1.0.0). User can freely access the online App to set up config parameters (e.g., domain size, substrates diffusion coefficients, custom data etc), simulate, and then plot simulation results. Interested readers can interactively run the model in a web browser at: https://nanohub.org/resources/pc4nanobio.

## 3. Results

After we developed the ABM-based nanotherapy model, we wished to explore how the model could be used to investigate interaction dynamics between anticancer drug-loaded NPs and cancer cells. We started by investigating how NP pharmacokinetic design parameters (internalization rate, drug release rate and decay rate) impact tumor growth dynamics. Next, we used the framework to simulate tumor response with different therapeutic schedules (NP injection dose and frequency). Then, we focused the investigation on NP “inheritance” across cell generations. To ensure that our results can generalize, we investigated cytostatic drugs (those that slow cycle entry) and cytotoxic drugs (those that induce cell death).

### 3.1. Large-scale NP pharmacokinetic design space parameter exploration

In this exploration, a single NP dose was injected at the beginning of simulation. For the NP decay, we only considered intracellular NP decay. NP “inheritance” was also ignored in this section, so daughter cells receive zero nanoparticles from their parent after division. Fig. 8 shows that the model is sensitive to the NP internalization rate and drug release rate, while it is relatively insensitive to the intracellular NP decay rate, especially in the case of faster drug release, because NPs would release the majority of their drug before they decay. We observed that slow drug release may improve treatments compared to fast release. This is consistent with “adaptive therapy” theory [50], where *containment treatment* with a limited dose may control cancer growth better than *aggressive treatment*. To explore how intracellular and extracellular NP decay influence treatment results, we performed another 9-parameter settings exploration, where we varied both the intracellular and extracellular NP decay rates. Fig. 9 shows that increasing both decay rates would significantly reduce cancer treatment efficacy even when drug is immediately released. This occurs because tumor cells would endocytose fewer nanoparticles due to quick decay of extracellular nanoparticles in the microenvironment. See Fig. 10 for tumor population dynamics for the two different scenarios simulated in Fig. 9. The tumor population dynamics show similar tendencies after about 12 days in Fig. 10(b) because only one NP dose was injected at the beginning of simulation. The majority of NPs had already been cleared from the tissue (e.g., by the renal system), modeled here as decaying boundary conditions) after several days, leaving very few NPs in the microenvironment.

**Figure 8:**
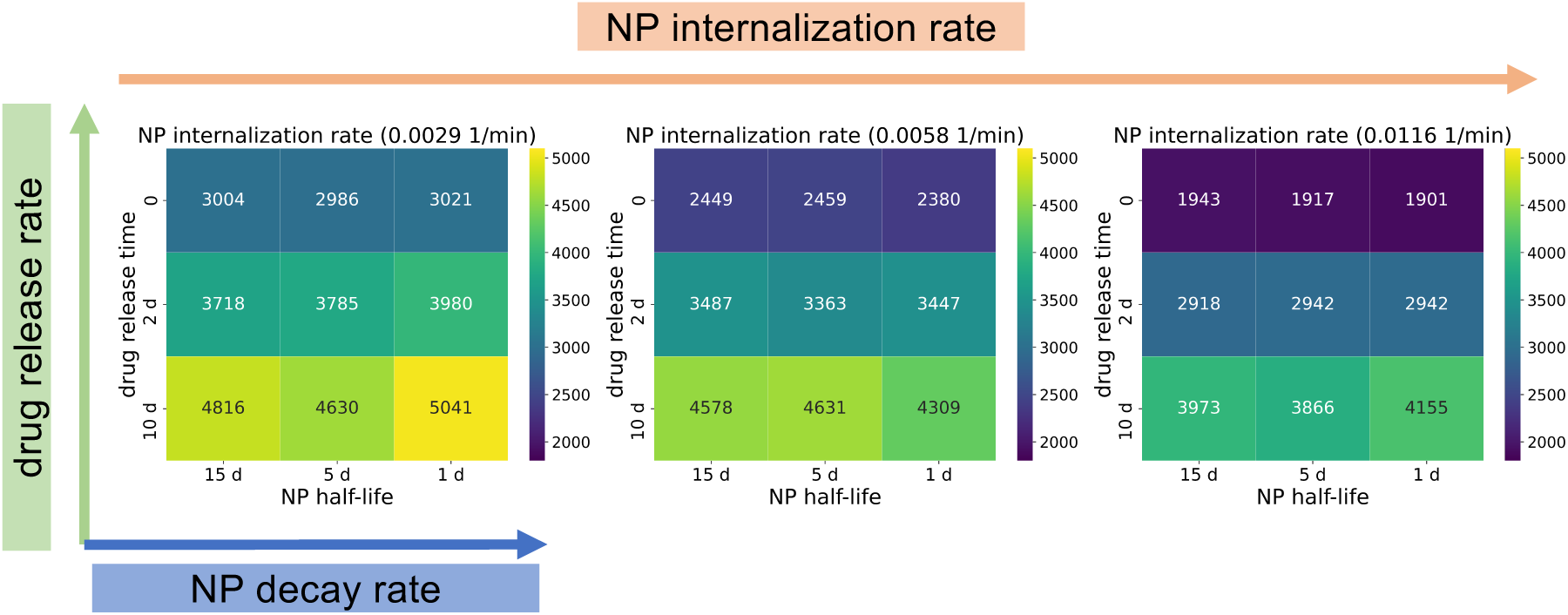
Simulation results of 27 parameter sets: heatmap of viable tumor cells population at 30 days with different NP design parameters (internalization rate, decay rate, drug release rate) (heatmap value is average of 10 runs for each parameter set). Dark blue squares denote the most effective treatments (fewest remaining tumor cells). We can observe that the model is sensitive to the internalization rate and drug release rate, while it is relatively insensitive to the intracellular decay rate, especially in the case of faster drug release, because NPs would release most of the loaded drug before their decay. See Fig. 9 for the exploration of intracellular and extracellular NP decay.

**Figure 9:**
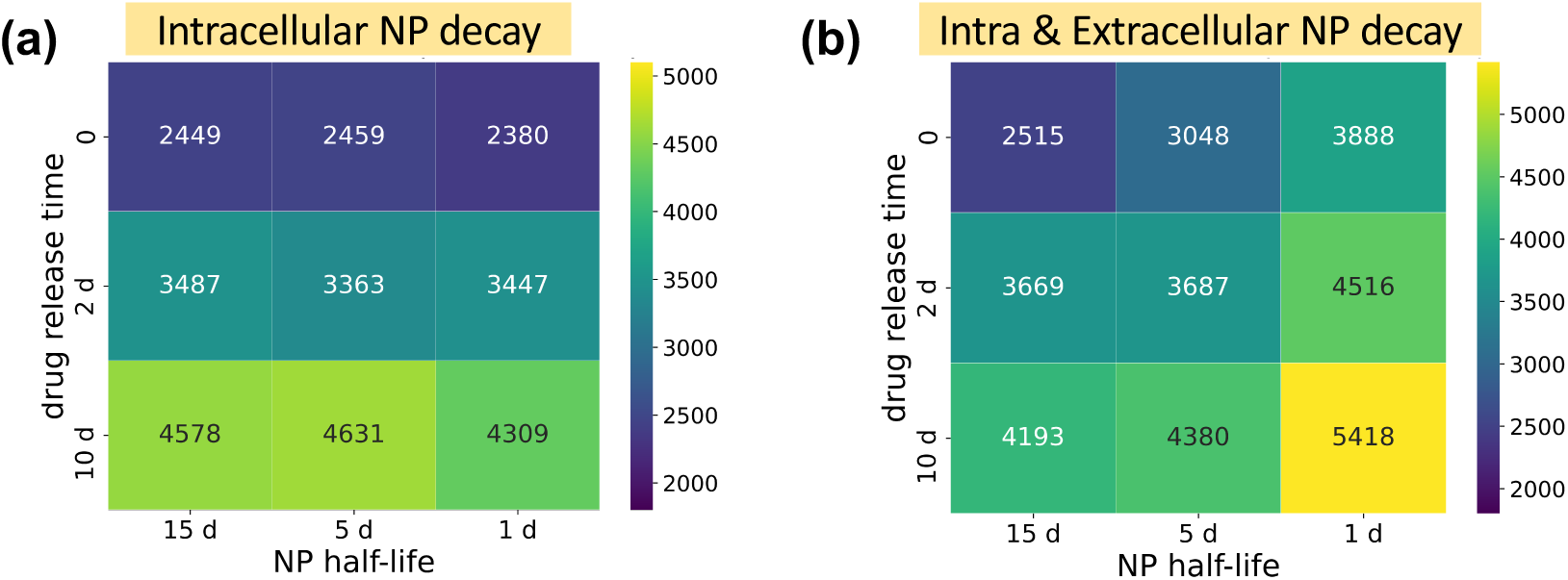
Heatmap of viable tumor cells population at 30 days in different NP decay scenarios (internalization rate is 0.0058 1/min). Dark blue squares denote most effective treatments (fewest remaining tumor cells). Treatment efficacy was most impacted by the drug release rate in all scenarios, but rapid extracellular NP decay could substantially reduce treatment efficacy, due to the reduction in NPs endocytosed by cells.

**Figure 10:**
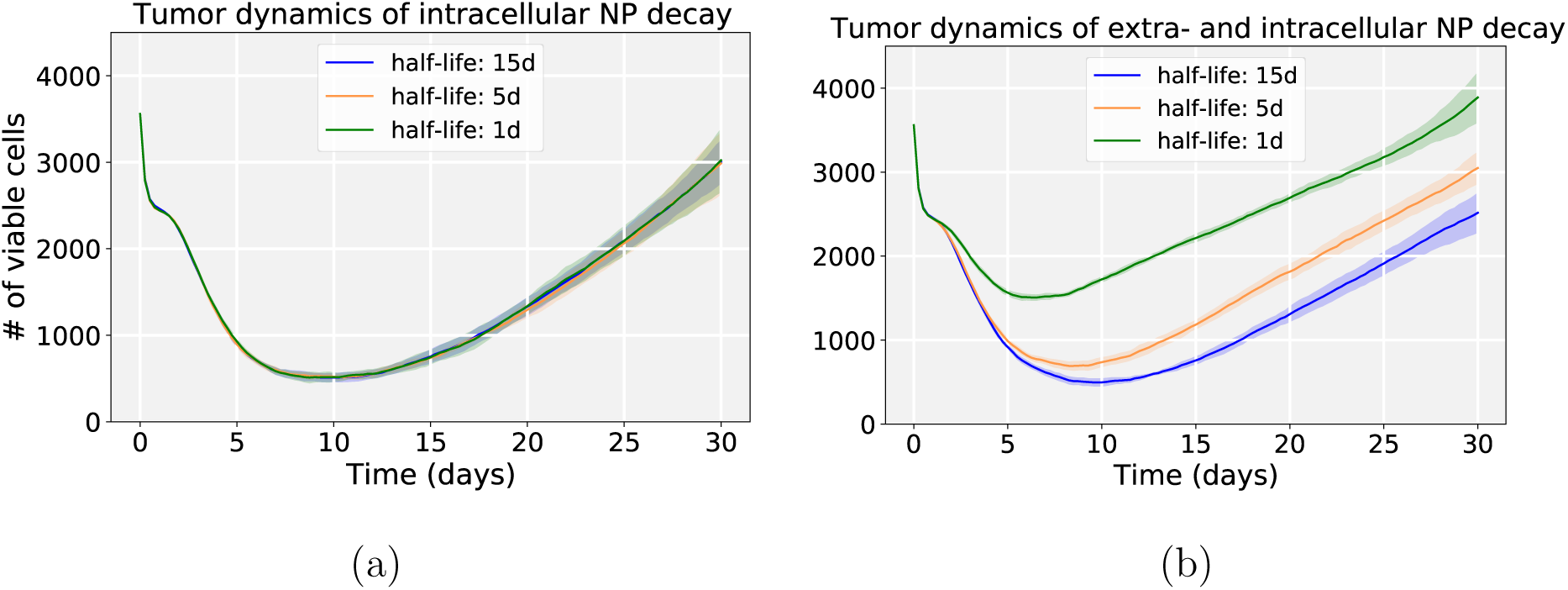
Tumor population dynamics of different NP decay rates for the simulations in Fig. 9 (when drug is *immediately released* by internalized NPs); (a) varying only intracellular NP decay; (b) varying both extra– and intracellular NP decay. Compared with only intracellular decay, increasing both decay rates could significantly reduce cancer treatment efficacy even when drug is immediately released. This is because tumor cells would endocytose fewer nanoparticles due to quick decay of extracellular nanoparticles in the microenvironment.

### 3.2. NP therapeutic schedules

We next explored NP therapeutic schedules with different NP injection doses and frequencies. From Fig. 11(a), we can find that multiple smaller doses may be better than single larger doses even though the total amount of injected NPs is the same. There are several factors that may contribute to this result. (1) From considering the boundary condition as a model of blood vessels connecting broadly the circulating, renal, and hepatic systems, larger doses do not necessarily ensure that all NPs would enter the tumor site from blood vessels, as some of the dose is cleared by the renal system. (2) Tumor cells need time to endocytose NPs from the microenvironment before they decay. (3) Due to slow NP diffusion in solid tumor tissue, a large dose may not penetrate the tumor from the periphery to the center in a short time, so tumor cells which are far away from periphery would not have access to sufficient NPs for a therapeutic response. (4) From an “adaptive therapy” perspective [50], higher doses may have decreased efficacy in containing tumor growth compared with lower doses. In Fig. 11(b), the second injection does not take effect immediately due to the time delay of NP internalization and drug release. Therefore, optimizing the NP pharmacokinetic design parameters and therapeutic schedule is vital for improving treatment. Fig. 12 gives simulation snapshots of four different therapeutic strategies.

**Figure 11:**
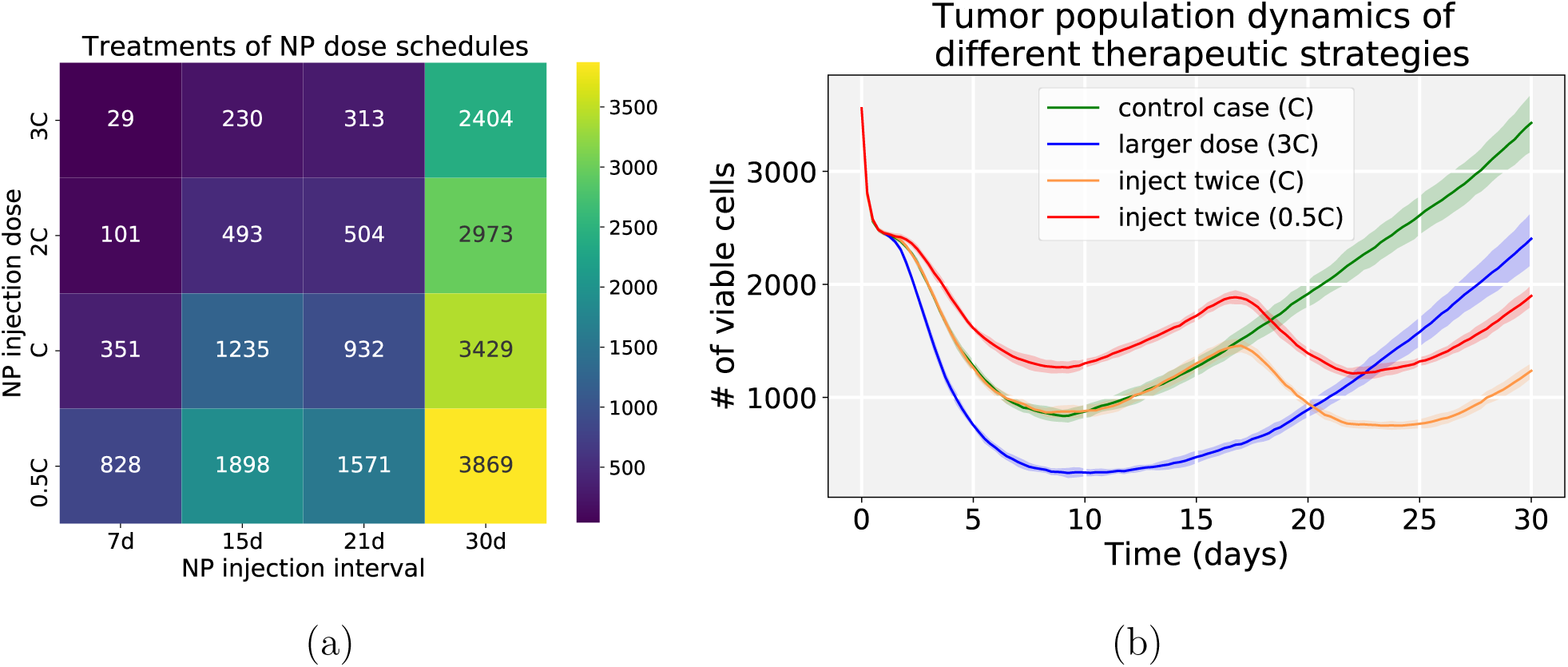
Simulation results of NP injection schedules. (a) Viable tumor cells population heatmap at day 30 (NP internalization rate: 0.0058 1/min; drug release time: 2 days; NP half-time: 5 days); (b) Tumor population dynamics of different therapeutic strategies, including a single regular dose (green), a single triple dose (blue), two regular doses (orange), and two half doses (red). Note that *C* is the default NP injection dose in the simulation. We can observe that multiple lower doses may have better performance that single larger doses to control the tumor (e.g., inject twice (0.5C) versus control case (C)).

**Figure 12:**
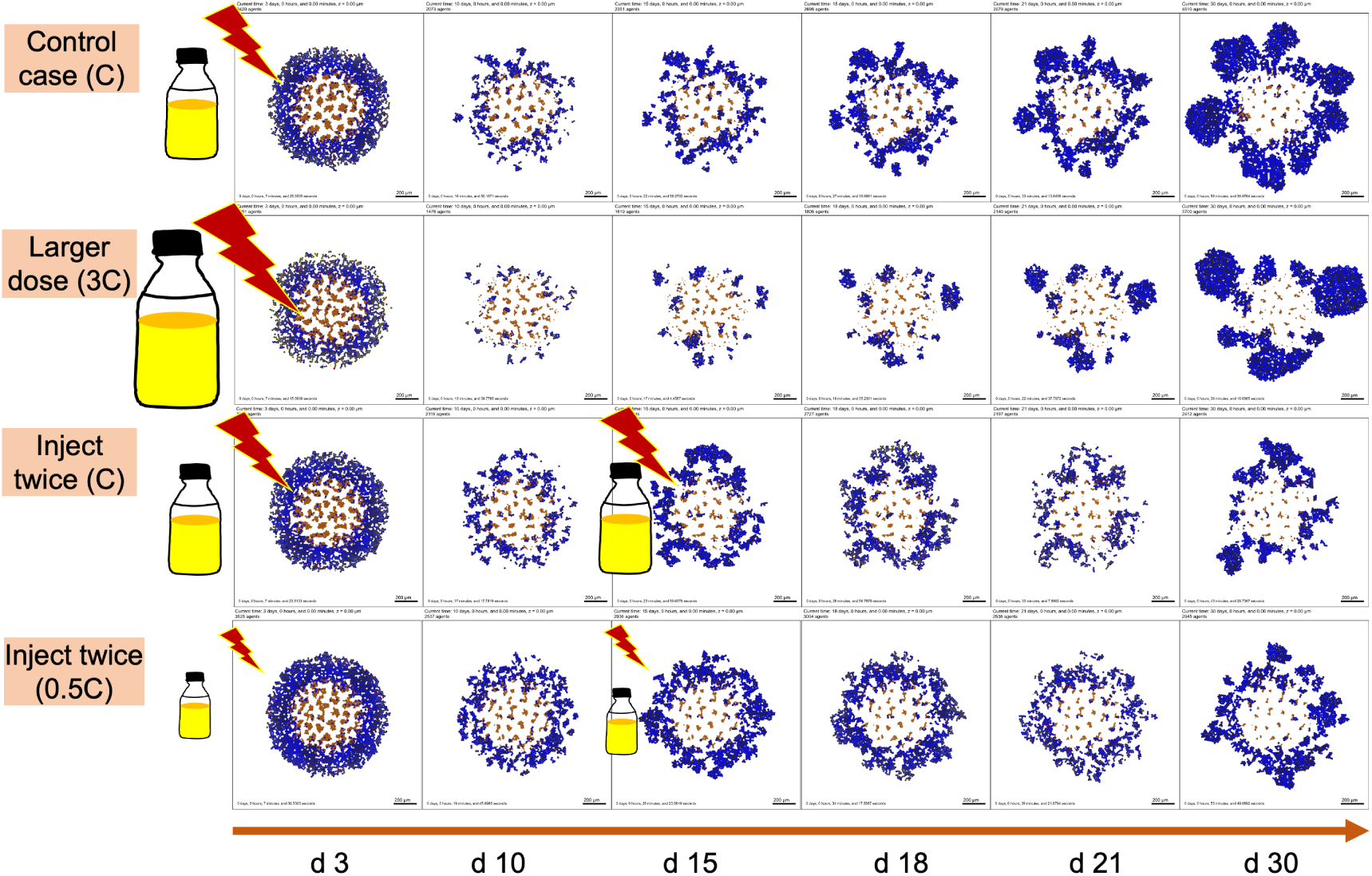
Snapshot of tumor patterns of different therapeutic strategies for the simulation in Fig. 11(b). **Legend**: blue (without drug effect), yellow (colored by drug effect), brown (necrotic), red (apoptotic). We can observe that the treatment of *inject twice* (0.5C) is better than *control case* (C) even the total dosage of two scenarios is the same. Additionally, *inject twice* (0.5C) works better than a single large dose (3C) because there are few NPs left in circulation at later stages for a single larger dose.

### 3.3. NP “inheritance” may improve cytotoxic chemotherapy

Experimental work has shown that NPs are potentially inherited by daughter cells during cell division [51, 52]; see Fig. 13 for experimental results of NPs inherited during mitotic process [51]. Lijster et al. recently used statistical modeling to investigate the impact of coefficient of variation (standard deviation over mean) of the number of nanoparticles per cell over the cell population for NP inheritance [53], finding that coefficient of variation is sensitive to the degree of asymmetry of nanoparticle inheritance.

**Figure 13:**
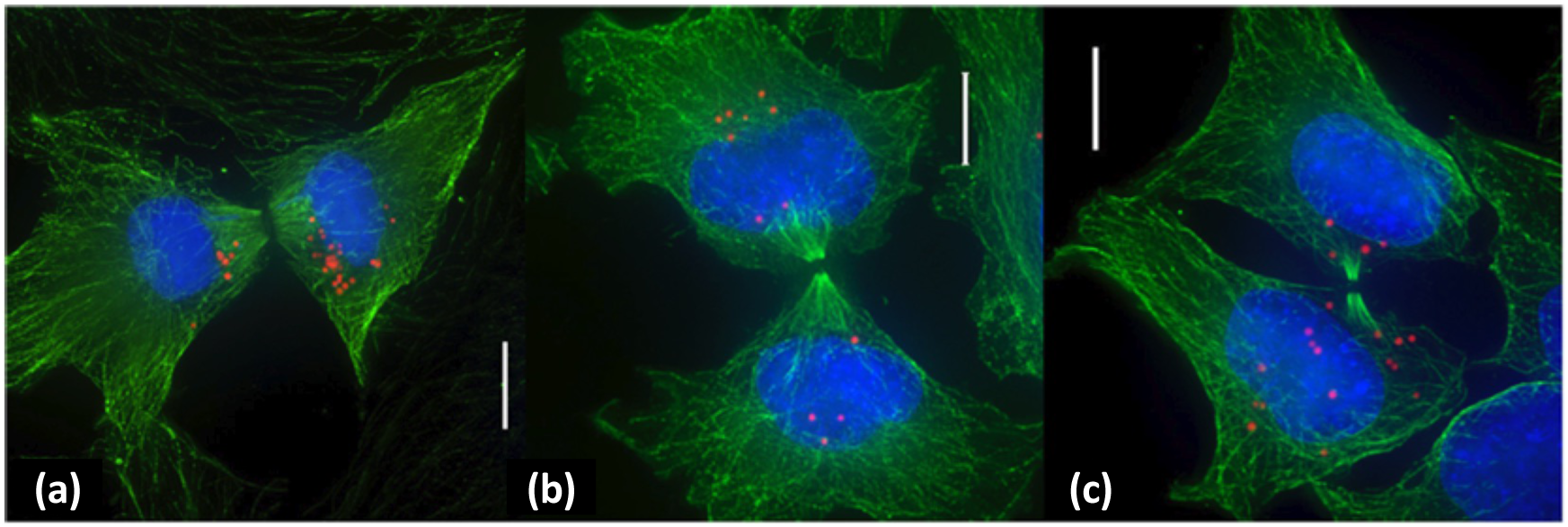
NP inheritance at cell division: Representative deconvolution microscopy images of cells undergoing cytokinesis. Microtubules are stained green, nuclei are labeled blue, and particles are shown in red. Scale bars: 10 *µm*. Adapted with permission from [51]. This is an unofficial adaptation of an article that appeared in an ACS publication. ACS has not endorsed the content of this adaptation or the context of its use.

To the best of our knowledge, no mechanistic modeling study has explored cancer nanotherapies where NPs can be “inherited” at cell division. Fig. 14(a-b) presents hypothesis simulation results of cellular NP internalization (with and without NP inheritance). From Fig. 14(c), we can observe that more tumor cells contain internalized NPs after 2.5 days of simulation if daughter cells inherit NPs at cell division, which raises the possibility of multi-generation treatments during nanotherapy. In this section, we explore how this “inheritance” would affect treatment response, in particular for cytostatic and cytotoxic anti-cancer drugs.

**Figure 14:**
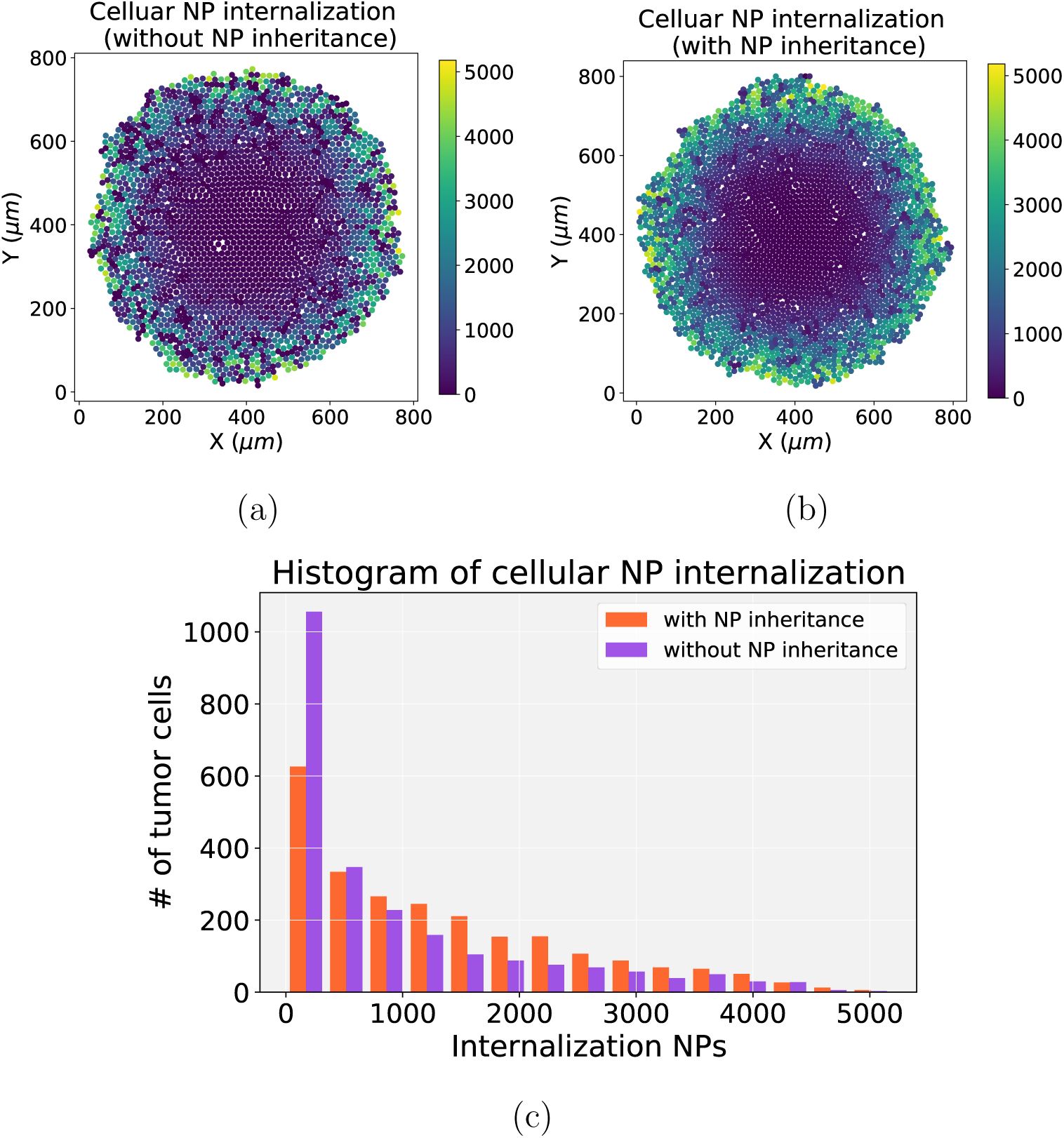
NP inheritance visualization. (a) Cellular NP internalization (without NP inheritance) after 2.5 days of simulation; (b) Cellular NP internalization (with NP inheritance); (c) Comparison of histograms of cellular NP of internalization in (a) and (b). We find that tumor cells would contain more NPs if NPs can be inherited at cell division, which raises the possibility of multi-generation treatments with nanotherapy. (Note that there are only cycling for tumor cells in (a) and (b)).

We first investigated the scenario of injecting a regular dose (*C*) once at *t* = 0. In Fig. 15, we find that cytotoxic treatment is moderately improved as more NPs are inherited, while there is no clear improvement for cytostatic chemotherapy. Cytostatic drugs inhibit cell cycling, making it difficult to transfer NPs to daughter cells. For cytotoxic drugs, the apoptosis rate is increased, but it has no effect on the cell cycling rate. Thus, when surviving cells can pass on their NPs to daughter cells at division, allowing the therapy to continue and cause additional apoptosis events that slow tumor growth. See Fig. 16.

**Figure 15:**
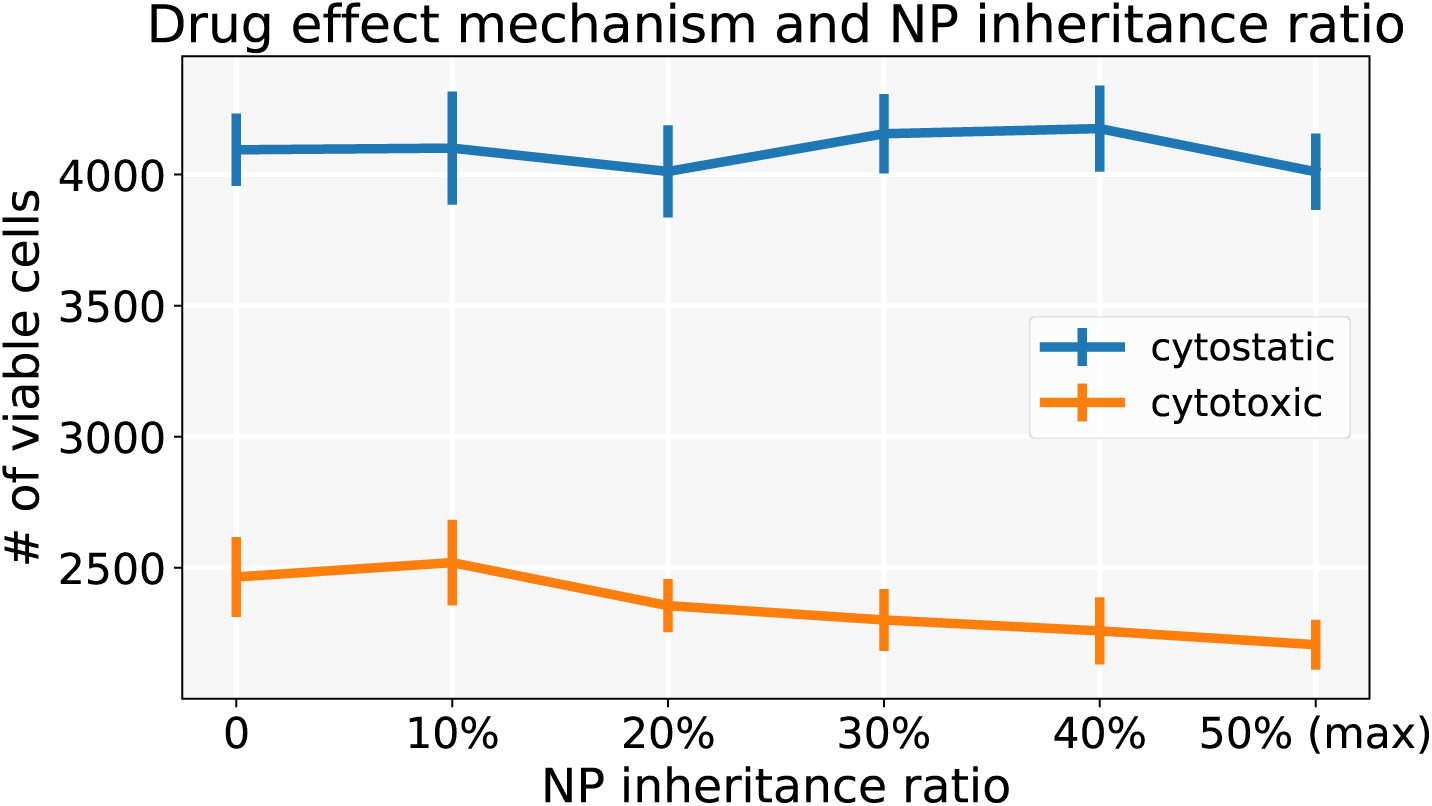
Comparison of cytostastic and cytotoxic drugs treatments with different NP “inheritance” ratio (**inject once** (*C*)). Note that 0 means daughter cells receive zero NPs from parent cell, and 50% means that each daughter cell gets 50% NPs (maximum). ErrorBar represents one standard deviation of 10 runs. We can observe that the cytotoxic treatment improves as more NPs are inherited, while there is no clear improvement for cytostatic chemotherapy.

**Figure 16:**
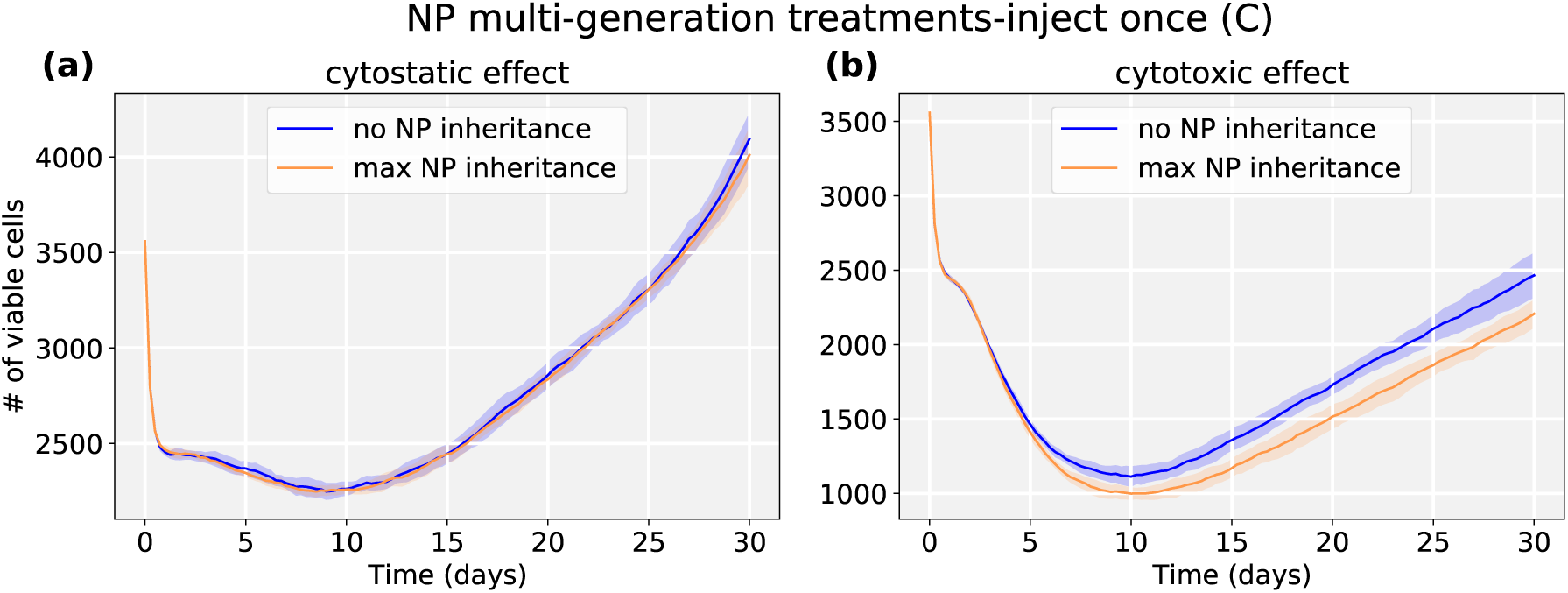
Tumor dynamics of cytostastic and cytotoxic effect treatments at 0 and maximum NP “inheritance” (**inject once** (*C*)). Cytostatic drugs inhibit cell cycling, making it difficult to transfer NPs to daughter cells. For cytotoxic drugs, the apoptosis rate is increased, but without an effect on the cell cycling rate, daughter cells can still receive inherited NPs from their parent cell before entering apoptosis.

### 3.4. Cytostatic chemotherapy may be improved by NP “inheritance” when cell division is not completely inhibited

We explored another scenario: injecting a half dose twice (0.5*C*, 0.5*C*) at *t* = 0, 15 days respectively. In this case, we found that *both* chemotherapies have better response if NPs are allowed to be inherited at cell division. See Fig. 17 and Fig. 18. Because smaller cytostatic drug doses cannot inhibit cell division in a short time (before entering into cycling phase) due to delay of NP internalization, some tumor cells still divide and transfer NPs to daughter cells.

**Figure 17:**
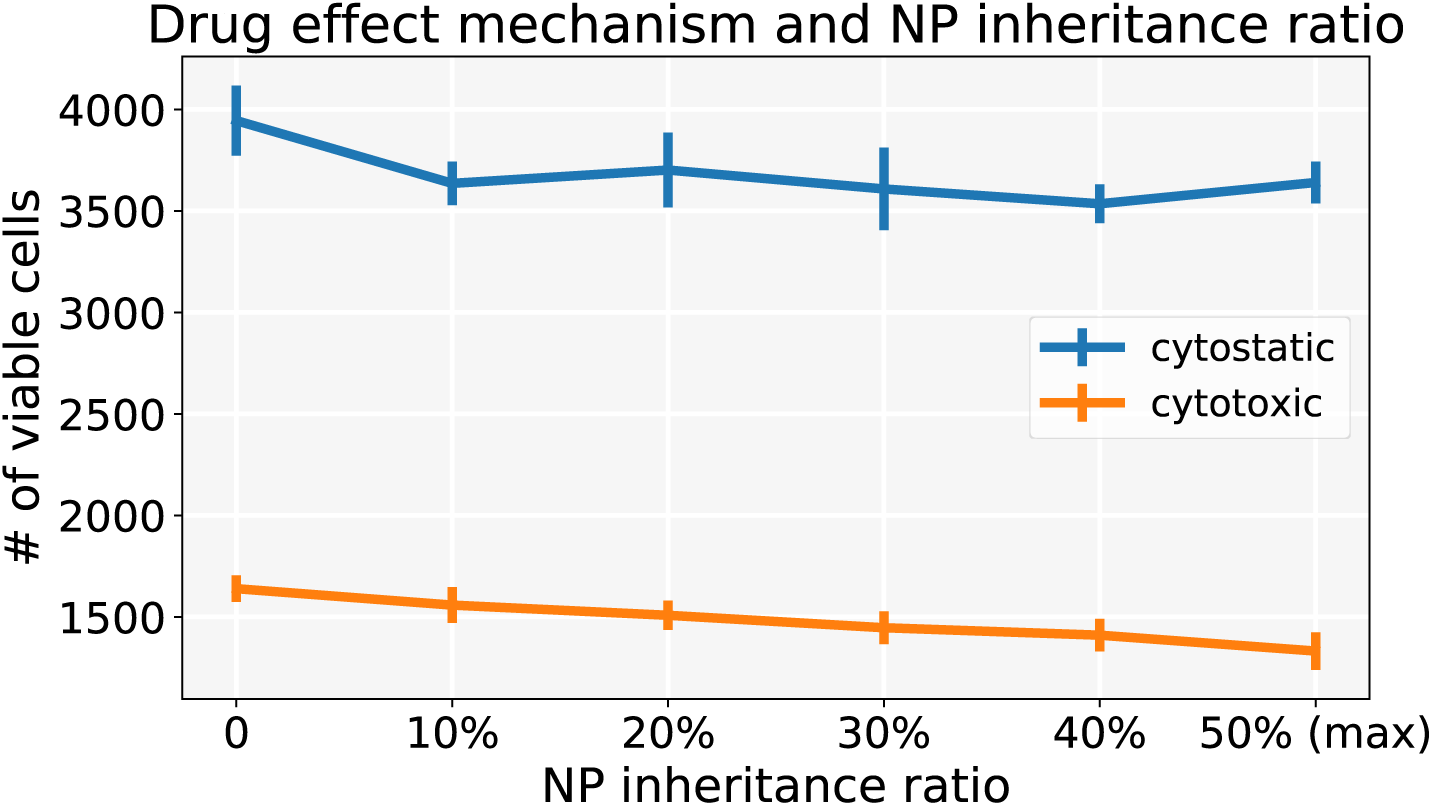
Comparison of cytostastic and cytotoxic drugs treatments with different NP “inheritance” ratio (**inject twice** (0.5*C*, 0.5*C*)). Error bar represents one standard deviation of 10 runs. Compared with Fig. 15, we can observe *both* chemotherapies have better response if NPs are allowed to be inherited at cell division under twice injections.

**Figure 18:**
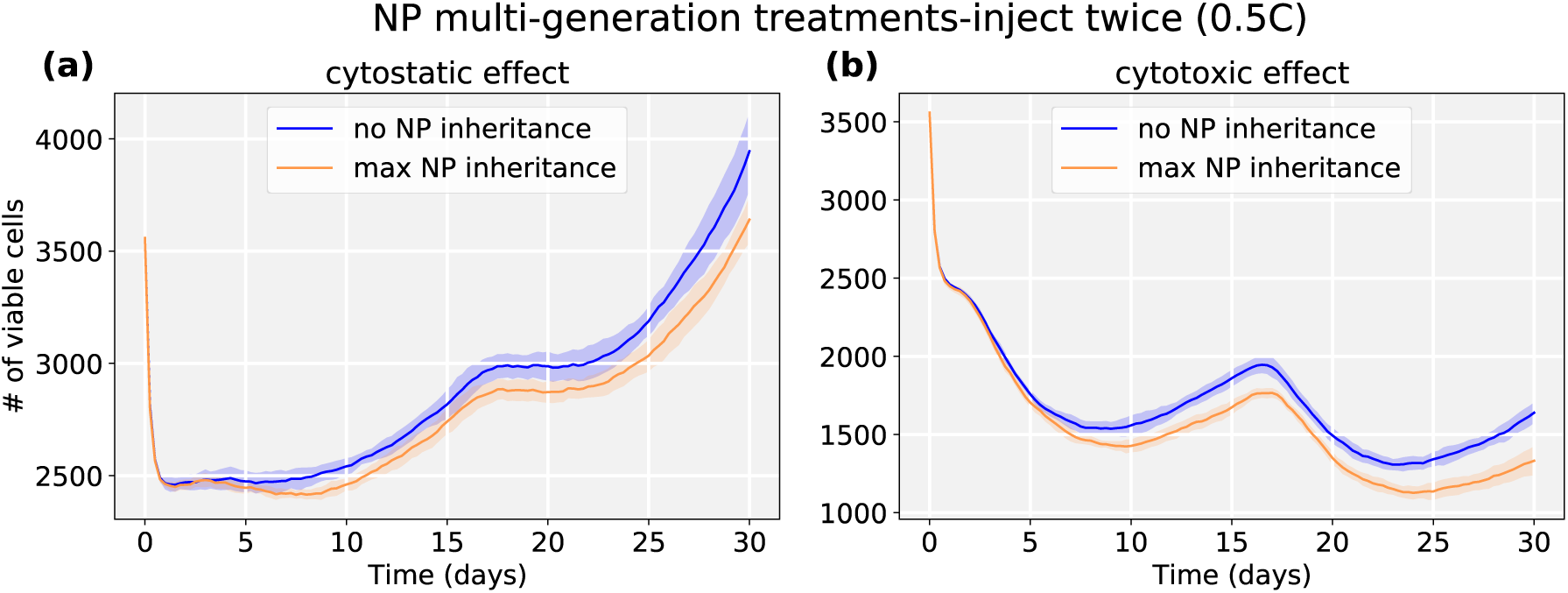
Tumor dynamics of cytostastic and cytotoxic effect treatments at 0 and 50% NP (maximum) “inheritance” (**inject twice** (0.5*C*, 0.5*C*)). We observe that two smaller doses lead to better treatments for both classes of anticancer drugs. Because smaller cytostatic drug doses cannot fully inhibit cell division in a short time (before entering into cycling phase) due to delay of NP internalization, some tumor cells still divide and transfer NPs to daughter cells.

## 4. Discussion

NPs offer attractive features for delivery of therapeutic agents to tumor cells, such as improved bio-distribution, protecting therapeutic agents from fast degradation in harsh microenvironments, improved binding rate via functionalized ligands that uniquely interact with receptors on tumor cell membranes, controlled intracellular drug release (via external stimuli, e.g., pH, enzymes, temperature), and enabling cancer immunotherapy through development of synthetic vaccines (e.g., incorporating DNA, siRNA, mRNA and protein) [3, 4, 9]. In this study, we developed a multicellular framework to evaluate cancer nanotherapy at the single-cell level. The ABM framework includes simulation modules of NP internalization (tracking how many NPs have been endocytosed by each cell); drug release (tracking the drug-release states for each cell as well as how much total drug is retained in individual cells; see Fig. 4); NP “inheritance” at cell division (see Fig. 14); and pharmacodynamic effects of anti-cancer drug effects on tumor cell phenotype (e.g., cycling, apoptosis).

In the exploration of pharmacokinetic design parameters, including NP internalization rate, NP decay rate, and NP drug release rate, we found that tumor response is sensitive to the NP internalization rate and drug release rate, while being relatively insensitive to the intracellular NP decay rate, especially in the scenario of faster drug release, because NPs would release most of the loaded drug before they decay (see Fig. 8). The exploration found that slow drug release may improve treatment compared to fast release due to *containment treatment* controlling cancer growth better than *aggressive treatment* with limited dose (based on “adaptive therapy” theory [50]). Furthermore, we varied both intracellular and extracellular NP decay rates to explore how these decays influence treatment response. We observed that increasing both rates would significantly reduce cancer treatment efficacy even when drug is immediately released (see Fig. 9 and Fig. 10), because tumor cells would endocytose fewer NPs due to fast decay of extracellular NPs in the microenvironment.

From the exploration of therapeutic schedules modifying different NP injection doses and frequencies, we observed that multiple smaller dosing injections may lead to better treatment outcomes than single larger doses even though the total amount of injected NPs is the same (see Fig. 11 and Fig. 12). A closer examination of the simulaitons gives an explanation: because NPs diffuse slowly into the microenvironment and tumor, a single large bolus of NPs may not reach and treat interior tumor cells, whereas a series of smaller treatments can wait for outer (treated) cells to respond and die, so that interior tumor cells can now be exposed to and endocytose NPs from subsequent rounds of treatment. In addition, some of the NPs may decay before reaching cancer cells far away from the tumor periphery. From an “adaptive therapy” perspective [50], larger doses may have worse efficacy in containing growth compared to smaller doses due to resource completion among cancer cells (e.g., cancer cells far away from oxygen sources may not have insufficient oxygen to proliferate). Therefore, it may be beneficial to optimize dosing and frequency of cancer nanotherapy taking into account these considerations.

In summary, the proposed nanotherapy model provides a platform for exploring NP design parameters, dosing regimens, and how NP “inheritance” may impact treatment response. Future work may focus on the NP-receptor binding dynamics (e.g., similar as virion-ACE2 binding [40, 44] and mRNA lipid nanoparticles (LNP) and receptor binding [41]), and mRNA vaccine-loaded LNPs for cancer immunotherapy [41, 54, 55]. With approximate preclinical and clinical data, this cancer nanotherapy framework could be used to design and calibrate novel patient-specific cancer control strategies. If clinical trials demonstrate that drug-loaded NPs can be safely and effectively designed to control or eliminate tumors in individual patients, they could be a powerful for use in future cancer patient digital twins [56, 57].

## Acknowledgements

PM, RH, and YW acknowledge funding from the National Science Foundation (1720625). PM, RH, and HR were funded in part by the Jayne Koskinas Ted Giovanis Foundation for Health and Policy and the National Cancer Institute (1U01CA232137). HF and PM were previously funded in part by the National Cancer Institute (1R01CA180149). VJ was funded in part by the National Science Foundation (DMR-1753182). We also thank FutureSystems (http://futuresystems.org) at the IU Digital Science Center, and Big Red 3 supercomputer at Indiana University Bloomington for powering high-throughput computational work.

## Appendix A. Code availability

Source code for this project hosted on GitHub repository is made public at: https://github.com/MathCancer/PhysiCell-nanobio.

## Appendix B. Main parameters used in the model

Table B.1 gives the main parameters we used in the simulation. Unless mentioned otherwise, parameters are at default values for Physicell 1.7.1.

**Table B.1:** Main parameters used in the simulation.

| Parameter | Value |
| --- | --- |
| Computational domain size | 1600 $\mu m$ by 1600 $\mu m$ |
| Computational mesh element size | 20 $\mu m$ by 20 $\mu m$ by 20 $\mu m$ |
| Oxygen diffusion coefficient | 100,000 $\mu m^2/\text{min}$ |
| Oxygen decay rate | 0.1 1/min |
| Oxygen boundary condition | 38 mmHg |
| NP diffusion coefficient | 6 $\mu m^2/\text{min}$ ([58]) |
| NP decay rate | 1.6667e-05 1/min (Estimated) |
| NP clearance rate | 4.8135e-04 1/min |
| maximum cellular NP internalization | 6000 ([46]) |
| tumor initial radius | 500 $\mu m$ |
| tumor cell maximum proliferation rate | 0.00072 1/min |
| tumor cell maximum necrosis rate | 0.002778 1/min |
| tumor cell apoptosis rate | 5.3167e-5 1/min |

## Notes

### Competing Interest Statement

The authors have declared no competing interest.

### Summary of Updates

This includes clarifications, refinements, and corrections to the introduction, methods, results, and discussion. The figures have also been streamlined.

